# Stretch-injury promotes activation of microglia with enhanced phagocytic and synaptic stripping activities

**DOI:** 10.1101/2023.03.24.534058

**Authors:** Anthony Procès, Yeranddy A. Alpizar, Sophie Halliez, Bert Brône, Frédéric Saudou, Laurence Ris, Sylvain Gabriele

## Abstract

Microglial cells must act as the first line of defense of the central nervous system, but they can be exposed to various mechanical signals that may trigger their activation. While the impact of chemical signaling on brain cells has been studied in detail, our current understanding of the mechanical signaling in microglia is still limited. To address this challenge, we exposed microglial cells to a single mechanical stretch and compared their behavior to chemical activation by lipopolysaccharide treatment. Here we show that stretching microglial cells results in their activation, demonstrating a strong mechanosensitivity. Stretched microglial cells exhibited higher Iba1 protein levels, a denser actin cytoskeleton and migrated more persistently. In contrary to LPS-treated cells, stretched microglia maintain a robust secretory profile of chemokines and cytokines, except for TNF-α, highlighting the relevance of this model. Interestingly, a single stretch injury results in more compacted chromatin and DNA damage, suggesting possible long-term genomic instabilities in stretched microglia. Using neuronal networks in compartmentalized microfluidic chambers, we found that stretched microglial cells exhibit enhanced phagocytic and synaptic stripping activities. Altogether, our results propose that the immune potential of microglial cells can be unlocked by stretching events to maintain brain tissue homeostasis after mechanical injury.

## 1. Introduction

Forces are constantly applied on the human body during a lifetime. In some pathological contexts, a rapid load can be exerted on brain tissue leading to large internal stress that result in stretching and compressive forces applied on brain tissues.[1] The inflammatory response following mechanical insults is considered as a major secondary injury mechanism that can induce long-term consequences, such as an increased risk for patients to develop neurodegenerative disorders, chronic traumatic encephalopathy, and amyotrophic lateral sclerosis in later stages of life.[2] In this neuroinflammatory context, the role of glial cells is to produce inflammatory mediators, scavenge cellular debris and orchestrate neurorestorative processes to promote neurological recovery. Among glial cells, microglia are the immune resident cells of the brain and play important roles in antigen presentation[3], phagocytosis[4], programmed cell death[5], vessel patterning[6] and neuronal plasticity including synaptic pruning.[7] Under physiological conditions, microglia adopts a monitoring state, extend their processes to scan the microenvironment searching for potential threats, participate to the neuronal networks maintenance releasing a minimal amount of cytokines and chemokines, and eliminate cellular debris and unnecessary synapses through phagocytosis.[8] During traumatic events and inflammatory episodes, microglial cells adopt a reactive mode, allowing microglia to orchestrate the immune response against perturbations of central nervous system (CNS) homeostasis. Depending on the nature of changes in stimulus, microglia can adopt different activation states, which correspond to altered microglia morphology, gene expression and function. It has been shown that microglia react to many chemical factors, such as lipopolysaccharides (LPS) or interferon-gamma (IFN-γ), which can trigger a reactive state[9]. Surprisingly, it was shown that microglia can undergo an activated state for days, weeks and even years after the trauma.[10,11] However, despite recent advances on the understanding of the mechanosensitivity of microglial cells[12,13], the mechanisms and functional consequences of a mechanical activation of microglial cells remain unclear.

A better consideration of the mechanobiological aspects of traumatic brain injury (TBI) is therefore critical to move the field forward. In this study, we sought to investigate the roles of a mechanical injury on microglial cells. We simulated traumatic mechanical stress by applying an uniaxial stretch of 20% of strain to microglial cells cultured on deformable elastomer membranes. Mechanically-activated microglial cells were compared to LPS-activated BV2 cells to gain further insights in the mechanisms and functional consequences of a mechanical injury.

## 2. Results

### 2.1. Single stretch of microglial cells results in their activation

We simulated traumatic mechanical stress by applying an uniaxial stretch to BV2 cells which are a type of cortical microglial cell derived from C57/BL6 mice.[14] BV2 microglial cells were grown on polydimethylsiloxane (PDMS) chambers coated with a mix of laminin and poly-L-lysine to ensure a specific engagement of transmembrane integrins. After 24 hours in culture, BV2 cells were subjected to a single uniaxial stretch of 20% in less than a second **(Fig. 1A and Supplementary Movie S1)**. The behavior of mechanically activated microglial cells were compared to a chemical activation with lipopolysaccharides (LPS), which is a commonly used pro-inflammatory stimulus for microglia, both *in vitro* and *in vivo* [15–17]. LPS evoked higher pro-inflammatory gene expression and also increased several anti-inflammatory genes [18]. Even though LPS-treated cells may not reflect the entire spectrum of activated microglial cells following TBI, it allows to compare mechanically-activated microglial cells to a well-known pro-inflammatory scenario.

**Figure 1.**
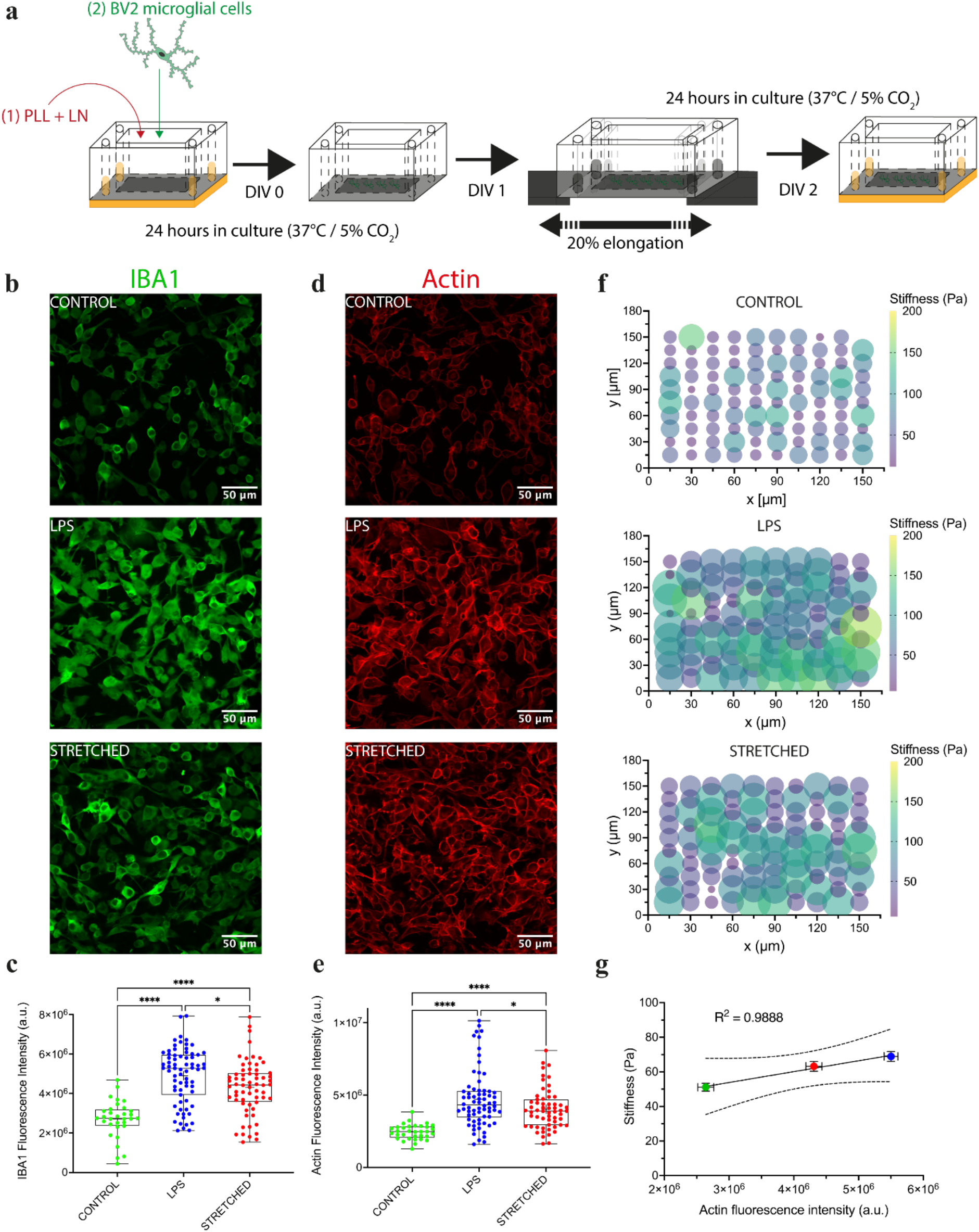
– Single stretch of microglial cells results in their activation. (A) BV2 cells were grown for 24 hours on deformable chambers coated with poly-L-lysine and laminin (PLL/LA). A deformable chamber was placed on an automatic stretcher and submitted to a uniaxial stretch of 20% of elongation in less than a second. Stretched chambers were then placed at 37°C and 5% CO_2_ for 24 hours. (B) Iba1 immunostaining of microglial cells without treatment (control), chemically-activated with LPS at 100 ng/ml for 24 hours (LPS) and mechanically-activated with a 20% stretch (stretched). (C) Total fluorescence intensity for Iba1 in control group (n=32), LPS (n=71), and stretched (n=64). (D) F-actin immunostaining of microglial cells for control, LPS and stretched cells (E) Total fluorescence intensity for Iba1 in control group (n=33), LPS (n=70), and stretched (n=59). (F) Cell stiffness probed with nanoindenter in function of (*x,y*) spatial coordinates for control (n=99), LPS (n=120) and stretched (n=109) groups. (G) Linear relation between stiffness of microglial cells in function of the actin fluorescence intensity (R^2^=0.9888). N=3 replicates for control and N=6 replicates for LPS and stretched groups. Scale bars are 50 μm. ns ≥ 0.05, 0.01 ≤ p^∗^ ≤ 0.05, 0.001 ≤ p^∗∗^ ≤ 0.01, p^∗∗∗^ ≤ 0.001.

We first assessed the state of activation of microglial cells by immunostaining the ionized calcium-binding adapter molecule (Iba1) that is specifically more expressed in activated microglia and macrophages[19]. Our findings showed that mechanically-activated and LPS-treated microglial cells exhibited an increased level of Iba1 protein fluorescence intensity compared to the control group **(Fig. 1 B-C)**, demonstrating that a 20% stretch injury can induce a reactive phenotype of microglial cells without any additional cell death **(Supplementary Fig. 1)**.

Knowing that Iba1 is an actin cross-linker, we hypothesized that the mechanical activation of microglia can induce the reorganization of their actin cytoskeleton. Interestingly, we showed that stretched and LPS-treated cells exhibited larger values of F-actin fluorescence intensity compared to the control group **(Fig. 1 D-E)**, suggesting that the mechanical activation of microglial cells leads to the strengthening of their actin cytoskeleton. We performed similar experiments on primary mouse microglial cells (PMCs) and we found a similar linear relation between fluorescence signals of Iba1 and actin **(Supplementary Fig. 2)**, validating our results obtained on BV2 microglial cells.

Based on the modulation of the actin cytoskeleton and its cross-linker Iba1 in stretch-injured microglia, we then characterize their elastic modulus with a nanoindenter working in liquid mode at 37 °C **(Fig. 1F and Supplementary Movie S2)**. Our findings indicated that control microglial cells were very soft with an elastic modulus of 51.1±2.2 Pa, in agreement with previous reports [20], while we observed a stiffening of LPS-treated and stretched microglia with an elastic modulus of 68.9±2.9 Pa and 63.2±2.9 Pa, respectively. Interestingly, we found a linear relation (R^2^=0.9888) between the cell stiffness and the actin fluorescence intensity **(Fig. 1 G)**.

### 2.2. Secretion of most pro-inflammatory chemokines and cytokines in microglial cells is not affected by stretch injury, except for TNF-α

Microglial activation results in the production of pro-inflammatory cytokines such as IL-1ß, IFN-γ, and TNF-α [21]. While release of these factors is typically intended to prevent further damages to CNS tissues, these cytokines may be toxic to neurons and glial cells [22]. Upon activation, microglia can undergo diverse states that display different cell surface receptors and intracellular markers, secrete different factors, and exhibit various functions. Knowing that LPS-treated and stretched microglial cells both adopt an activated state, we then characterized their secretory profile to get more insights into the phenotype associated with a mechanical injury.

We performed Mesoscale Discovery (MSD) electrochemiluminescence multiplex immunoassays to analyze the concentration of IFN-γ, IL-1ß, IL-2, IL-5, IL-6, IL-10, IL-12p70, KC/GRO, and TNF-α in the supernatant of untreated, LPS-treated and mechanically-activated microglial cells. As shown in **Fig. 2A**, our results indicated that 7 cytokines (IFN-γ, IL-2, IL-6, IL-10, IL-12p70, KC/GRO, and TNF-α) were significantly more secreted by LPS-activated cells compared to untreated microglia, whereas IFN-γ (**Fig. 2B)** and IL-1ß remained unaffected. It is worth to notice that KC/GRO concentration has tendency to be increased in stretched microglial cell media **(Fig. 2C),** while it is statistically not significant (p=0.4675). We found that the concentration of TNF-α increased in the culture medium of LPS-treated (28732±1149 pg/ml) and to a lesser extent in mechanically-activated microglia (4025±1283 pg/ml) compared to healthy microglia (2590±953 pg/ml) **(Fig. 2D)**. In contrary to LPS-treated cells, these results show that mechanically-activated microglia maintain a robust secretory profile of chemokines and cytokines, except for TNF-α, highlighting the relevance of this model. The secretion of cytokines by activated microglial cells can be different regarding the trigger for cell activation [18] emphasizing the multiple facets of activated microglial cells. Previously, we have shown that TNF-α secreted by stretched astrocytes is a key player of the synaptic loss in neuronal networks, suggesting a complex interplay between mechanically injured glial cells [23].

**Figure 2.**
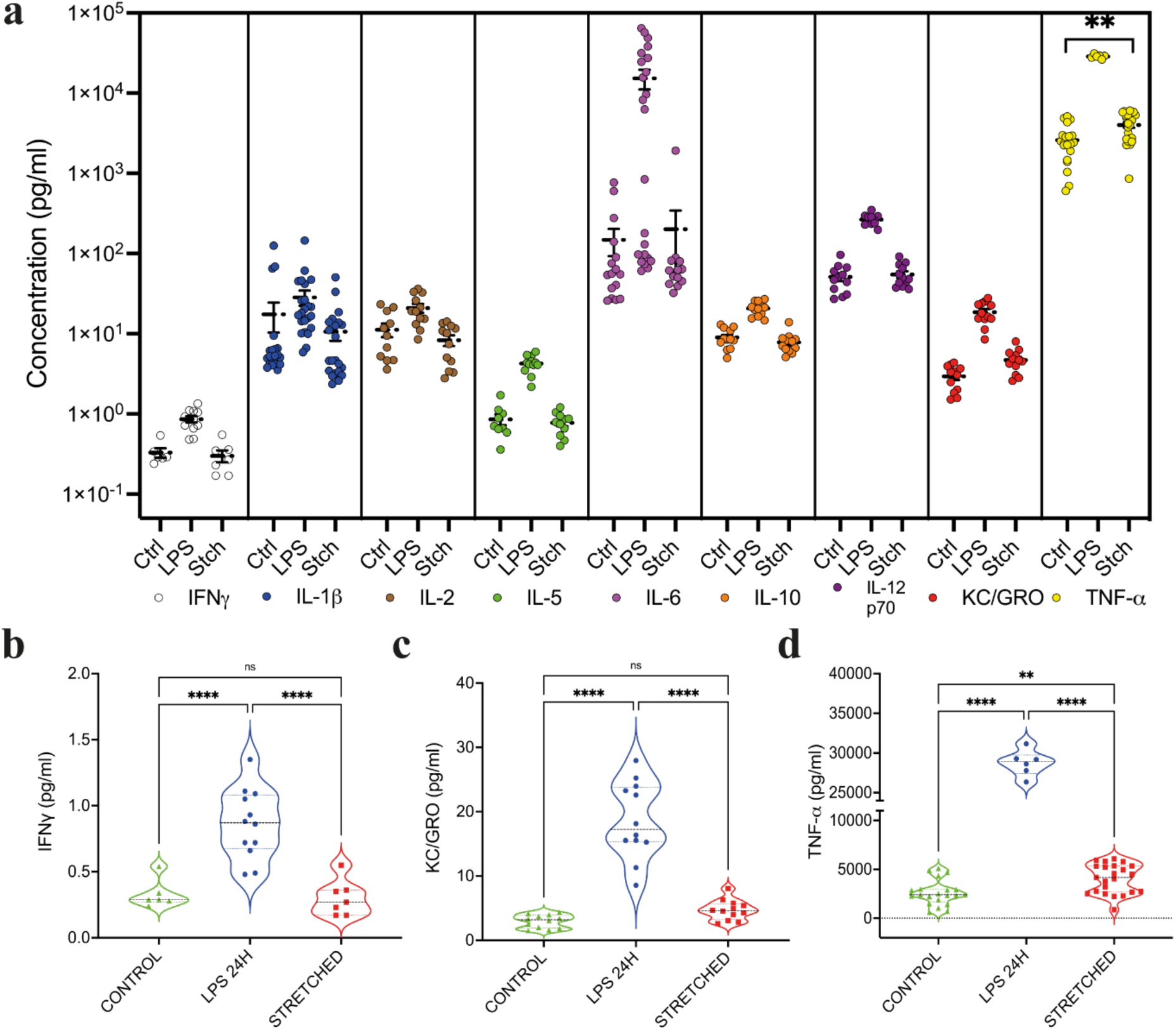
– Secretion of most pro-inflammatory chemokines and cytokines in microglial cells is not affected by a stretch injury, except for TNF-α. (A) Nested plots showing the concentration of nine cyto-chemokines (IFN-γ, IL-1β, IL-2, IL-5, IL-6, IL-10, IL-12p70, KC/GRO, and TNF-α) in the media of healthy, LPS-treated and mechanically-activated microglial cells. (B) IFN-γ concentration (control: n=6; LPS: n=6; stretched: n=6). (C) KC-GRO concentration (control: n=6; LPS: n=6; stretched: n=6). (D) TNF-α concentration (control: n=11; LPS: n=6; stretched: n=12). ns is not significant, 0.01 ≤ p^∗^ ≤ 0.05, 0.001 ≤ p^∗∗^ ≤ 0.01, p^∗∗∗^ ≤ 0.001.

### 2.3. Mechanical activation leads to a more persistent migration

While accumulating evidence suggests that the transition between a surveilling and a reactive mode in microglia induces a modulation of their morphology and migratory behavior[9], the impact of a mechanical injury on both properties of microglial cells is still unclear. To address this question, we assessed morphological and dynamic characteristics of BV2 cells with time-lapse microscopy experiments by using CellTracker, a live fluorescent dye. Our findings showed that BV2 cells adopted a larger perimeter (100.9±36.7 µm) and spreading area (500.1±180.1 µm^2^) in response to LPS treatments. Moreover, confocal imaging revealed that LPS-treated BV2 cells exhibited a larger cellular volume (7049±3342 µm^3^) than control cells (2080±1140 µm^3^). Our findings indicated that mechanically-activated BV2 cells were characterized by a larger spreading area (407.4±179 µm^2^, **Fig. 3A**) and a larger perimeter (92.4±40.4 µm, **Fig. 3B**) than control cells. However, we did not observe any significant modifications of the cellular volume (3101±1491 µm^3^) in mechanically-activated cells (**Fig. 3C**).

**Figure 3.**
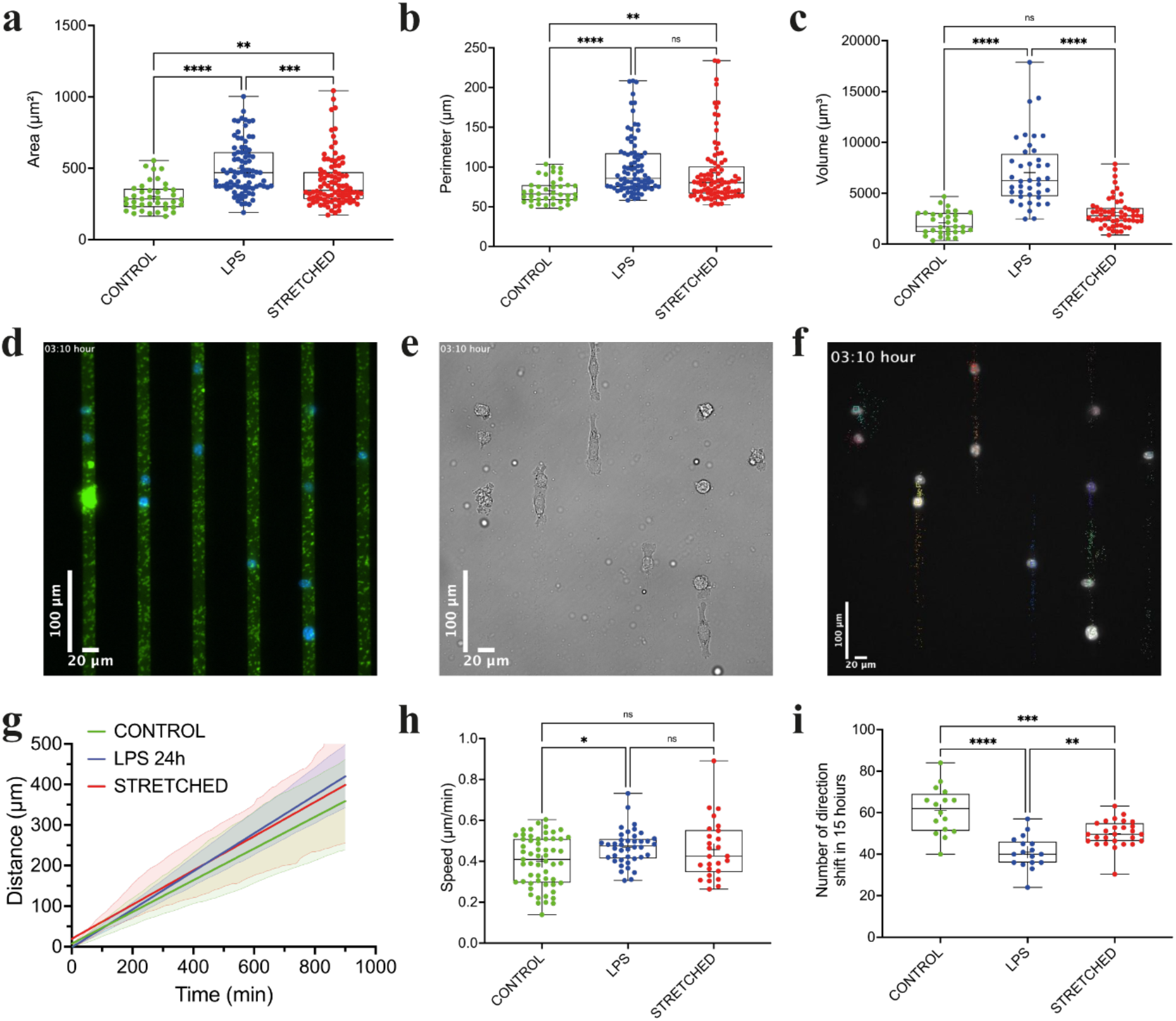
– Mechanically-activated microglial cells are larger and more persistent. (A) Spreading area, (B) perimeter and (C) cellular volume for control (n=39, in green), LPS-treated cells (n=85, in blue) and stretched cells (n=91, in red). Typical images of microglial cells migrating (D) on PLL-LA microstripes of 15 μm width (green) and (E) observed in Differential Interferential Contrast (DIC) mode. (F) The nucleus was stained with DAPI (in blue) to perform an automatic tracking on time-lapse experiments of 15 hours with a frame rate of 10 minutes. Temporal evolution of the travelled distance and (H) average migration speed for control (n=58, in green), LPS-treated cells (n=40, in blue) and stretched cells (n=26, in red). (I) Number of direction shifts for control (n=16, in green), LPS-treated cells (n=17, in blue) and stretched cells (n=26, in red). Scale bars are 20 μm. ns is not significant, 0.01 ≤ p^∗^ ≤ 0.05, 0.001 ≤ p^∗∗^ ≤ 0.01, p^∗∗∗^ ≤ 0.001.

Considering that a stretch injury induces a strengthening of the actin cytoskeleton and a recruitment of Iba1, which interacts with RAC GTP-ases that participate in lamellipodial protrusion via the ARP2/3 complex [24], we aimed to probe the migratory behavior of mechanically-activated microglial cells. We studied the migration of BV2 microglial cells for 15 hours using protein microstripes of 15 µm wide to standardize our migration assays (**Supplementary Movie S3**). We used time-lapse microscopy with live fluorescent labelling of the nucleus **(Fig. 3D-E)** to track the cell displacement over time **(Fig. 3F and Supplementary Movie S4)**. Our results showed that control microglial cells have a mean velocity of 0.40±0.12 µm/min, whereas LPS-treated and mechanically-activated microglial cells were slightly faster (0.47±0.09 µm/min and 0.46±0.15 µm/min, respectively) **(Fig. 3G-H)**. In addition, our results showed that LPS-treated and mechanically-activated BV2 cells performed less direction changes (41±7.8 and 50±6.4, respectively) than control (61±11.4) cells **(Fig. 3I)**. These results suggest that activated microglial cells adopt a more persistent mode of migration, with less back-and-forth movements, leading to more efficient goal-directed migration (**Supplementary Fig. 4**).

### 2.4. Stretch injury results in chromatin compaction and DNA damage

During their migration within in the cerebral parenchyma, microglial cells undergo important deformations to squeeze in narrow spaces. As the largest and stiffest organelle in eukaryotic cells [25], the nucleus is constantly subjected to intrinsic and extrinsic forces that can lead to various nuclear deformations [26], which also contributes to cellular perception of mechanical stimuli [27,28]. The nucleus must be considered not only as the primary site of gene replication and transcription but also as a fundamental mechanotransduction component of the cell, capable of mechanosensing and orchestrating key cellular functions in response to mechanical stimulation [26].

To understand whether a single stretch of 20% can affect the nuclear integrity of microglial cells, we first immunostained the nucleus with diamidino-2-phenylindole (DAPI), which selectively bind to the minor groove of double-stranded DNA. Using 3D confocal images, we found that nuclear volume of mechanically-activated microglial cells (957±274 µm^3^) was larger than control (424.5±109.8 µm^3^) and LPS-treated (543.3±195.8 µm^3^) cells **(Supplementary Fig. 3)**, suggesting that mechanical activation of microglia leads to elevated nuclear influx accompanied by nuclear volume expansion.

We then assessed whether the influx of cytoplasmic constituents could affect condensation state of the chromatin [29,30]. As shown in **Fig. 4A-B**, our results indicated that mechanically-activated microglial cells showed larger domains of chromatin compaction (0.15±0.04 a.u.) than control (0.07±0.03 a.u.) and LPS-treated (0.08±0.02 a.u.) cells. Interestingly, these results indicated that a single stretch of 20% induced more condensed chromatin states and thus can potentially affect gene expression, while the chromatin organization is not affected by a chemical activation with LPS treatment. To go a step further, we used immunocytochemical assays to study the potential presence of the phosphorylated form of γH2Ax (**Fig. 4C**), that results from double-strand breaks [31]. As shown in **Fig. 4D**, we found a substantial increase of the number of γH2Ax foci in mechanically-activated microglia nuclei (25.9±20 foci), indicating that many DNA breaks occur in response to a stretch injury (**Supplementary Movie S5**). The number of foci was statistically similar between the control group (10.8±7.6 foci) and the LPS group (14.2±10.4 foci) **(Fig. 4D)**.

**Figure 4.**
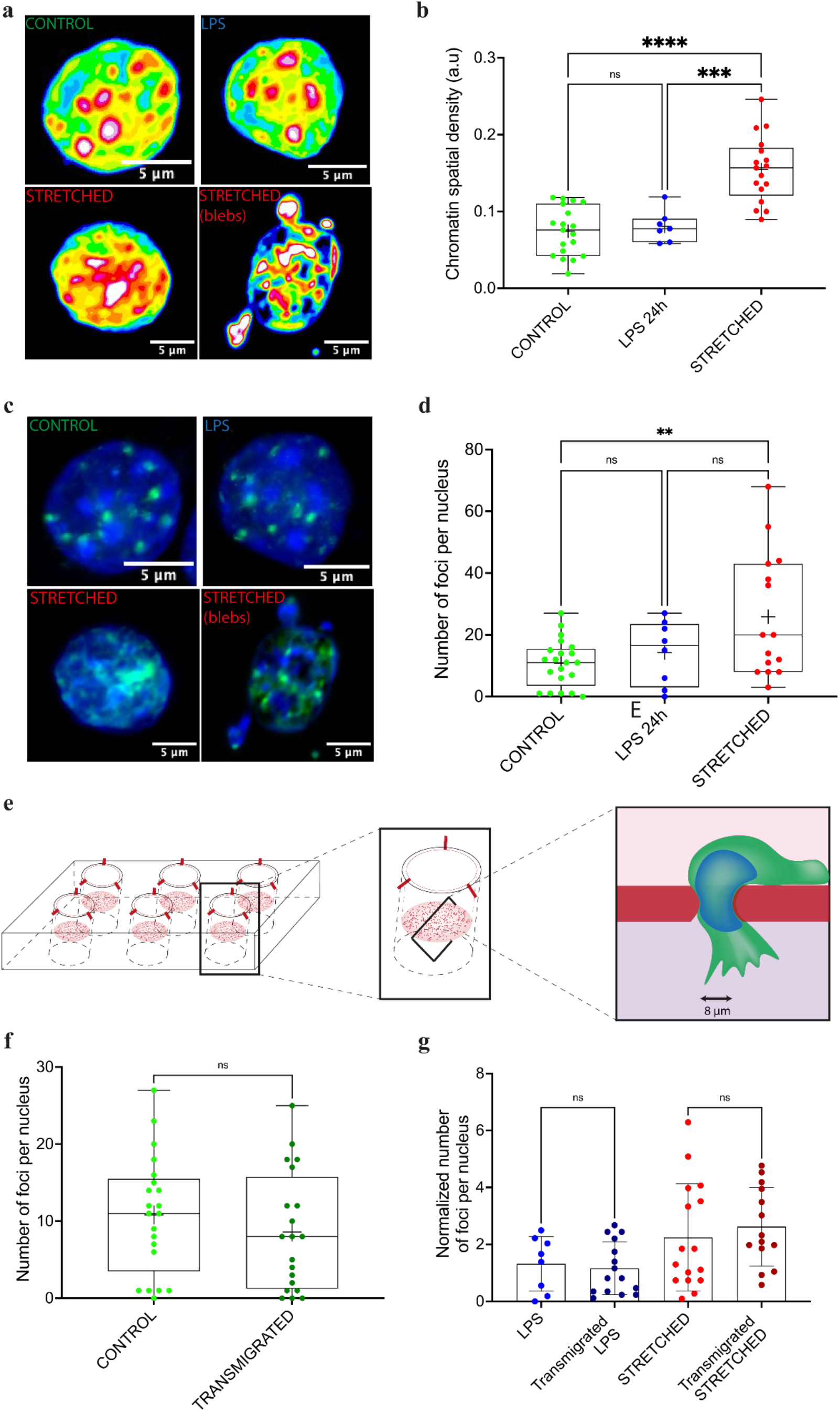
– Stretch injury results in more compacted chromatin and DNA damage, whereas confined transmigration does not. (A) Typical confocal images of nuclei with the fluorescence intensity of DAPI digitized (0-255 bits) and color coded (from high to low: white, purple, red, orange, yellow, green, light blue and dark blue). Highly condensed domains show higher fluorescence intensity with respect to the less condensed ones. (B) Average chromatin spatial densities of control (n=19, in blue), LPS-treated cells (n=7, in green) and stretched cells (n=17, in red). (C) Typical confocal images of immunostained nuclei for DAPI (in blue) and γH2Ax foci (in green) (D) Number of foci per nucleus for control (n=23, in blue), LPS-treated cells (n=7, in green) and stretched cells (n=17, in red). (E) Schematic representation of a transwell migration assay. Microglial cells were seeded on a culture insert with a porous membrane of teflon (pores of 8 μm in diameter) placed inside a 6-well plate. The culture insert was filled with a serum-free medium (light pink) and a complete medium was placed in the well of the plate (dark pink). The serum gradient triggers the confined transmigration of microglial cells through the narrow pores. (F) Number of foci per nucleus of control (n=23, light green) and transmigrated control cells (n=7, dark green). (G) Normalized number of foci per nucleus control for control and transmigrated LPS-treated and stretched cells. ns is not significant, 0.01 ≤ p^∗^ ≤ 0.05, 0.001 ≤ p^∗∗^ ≤ 0.01, p^∗∗∗^ ≤ 0.001.

Based on these results, we then investigated the mechanical sensitivity of chemically and mechanically activated microglial cells to 3D confined migration. To this aim, we performed transmigration assays through a porous membrane with pores of 8 µm in diameter **(Fig. 4E)**. We did not observe any differences in the number of nuclear foci between transmigrated and control cells **(Fig. 4F)**, suggesting that a transmigration through a porous membrane does not induce additional DNA defects. Furthermore, LPS-treated or mechanically-activated microglial cells did not exhibit any additional γH2Ax foci after transmigration **(Fig. 4G)**. Altogether, these results demonstrated that transmigration through 3D confined spaces does not lead to severe nuclear deformations and thus DNA damages in contrary to a 20% stretch injury, suggesting different impacts on nuclear integrity between endogenous stress during confined migration and exogenous stress during tissue stretching.

### 2.5. Phagocytosis and synaptic stripping are enhanced in stretch injured microglia

As the resident macrophages of the brain tissues, microglia must assume antigen-presenting tasks, as well as clearing of cellular debris or pathogens threatening brain homeostasis. Phagocytosis describes the process by which a cell recognizes, engulfs and digests a target that is ≥ 1 µm in size, including dead or dying cells, during physiological and pathological conditions [32,33]. To assess the phagocytotic capacity of microglial cells, we introduced pre-opsonized fluorescent latex beads into the culture medium for 1 hour and then we washed the excess of beads to only count those that have been phagocytized using 3D confocal images **(Fig. 5A)**. We found a 3-fold increase of the phagocytic activity in mechanically-activated microglial cells (1.5±0.4 beads/cell), while we observed a slight augmentation in LPS-treated cells (0.9±0.5 bead/cell) compared to the control group (0.6±0.3 bead/cell) **(Fig. 5B)**.

**Figure 5.**
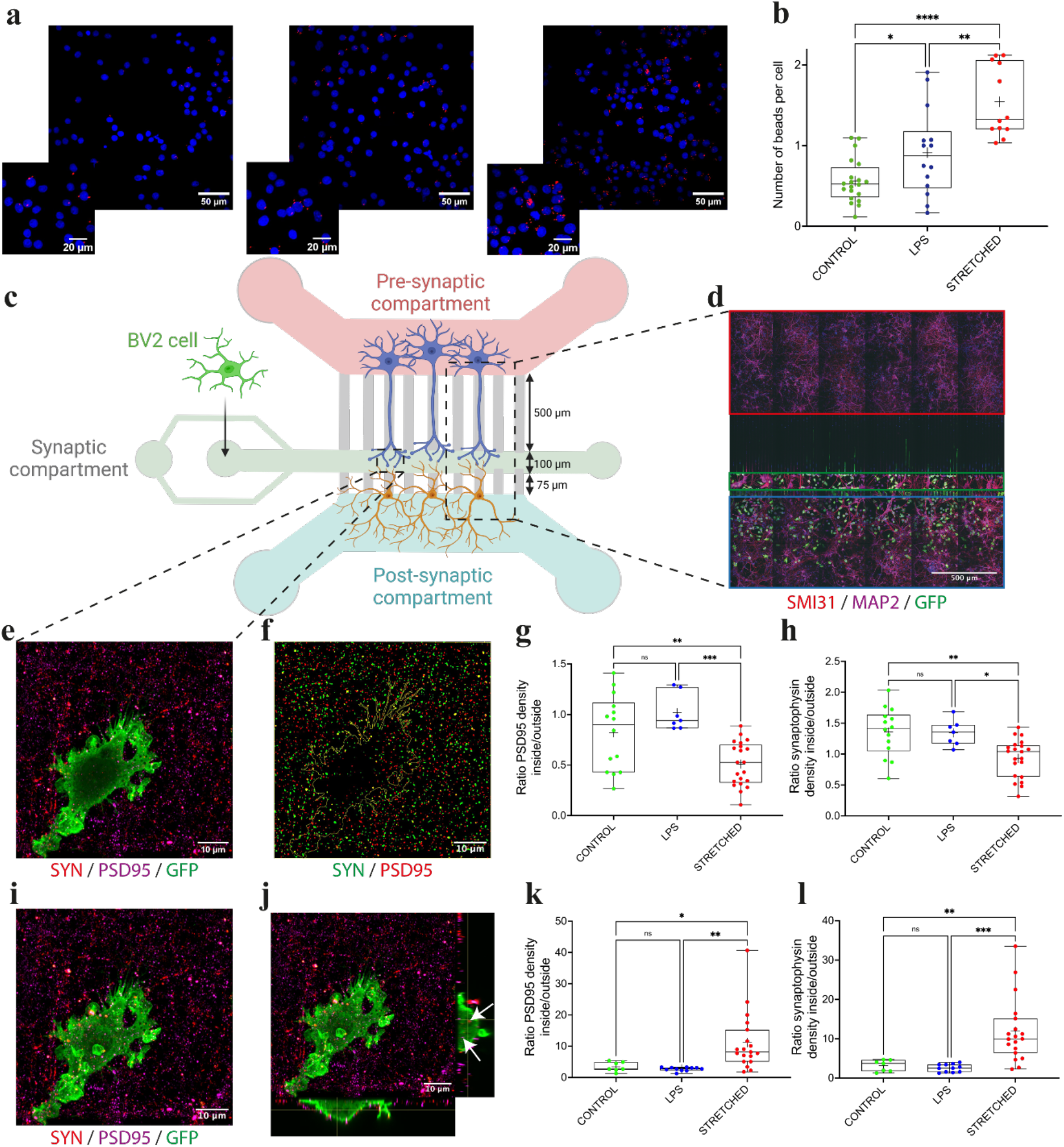
– Mechanical activation of microglial cells triggers increased phagocytosis and synaptic stripping activities. (A) Confocal images of nuclei (stained with DAPI in blue) and fluorescent latex beads (in red) in microglial cells. (B) Mean number of beads per cell for control (n=20, in green), LPS-treated (n=14, in blue) and stretched cells (n=12, in red), with N≥3 replicates. (C) Schematic representation of the microfluidic chamber. Primary cortical neurons were seeded in pre-synaptic (in light red) and post-synaptic (light blue) chambers. BV2 cells were seeded in the synaptic compartment (light green) at neuronal network DIV10. (D) Typical confocal image of the microfluidic chamber immunostained for SMI31 (red), MAP2 (purple) and GFP (green). GFP-BV2 microglial cells migrated to synapses and through lower and upper channels (3 μm wide). (E) Confocal image (3 μm-depth) of microglial cells immunostained for synaptophysin (red), PSD95 (purple) and GFP (green). (F) Processed image showing the microglial cell contour (yellow line) and pixelized dots for synaptophysin (green) and PSD95 (red). Ratio between inside and outside microglial cell densities for (G) PSD95 and synaptophysin for control (n=14, in green), LPS-treated (n=7, in blue) and stretched cells (n=21, in red). (I) Confocal image of a microglial cell immunostained for synaptophysin (red), PSD95 (purple) and GFP (green). (J) Orthogonal views of the confocal image showing the presence of synaptophysin (red) and PSD95 (purple) dots inside the cell (white arrows). Ratio between inside and outside microglial cell densities for (K) PSD95 and (L) synaptophysin for control (n=7), LPS (n=11) and stretched cells (n=19), without the first 3 μm of confocal acquisition. ns ≥ 0.05, 0.01 ≤ p^∗^ ≤ 0.05, 0.001 ≤ p^∗∗^ ≤ 0.01, p^∗∗∗^ ≤ 0.001.

Based on our previous results, we were wondering whether a mechanical activation of microglial cells could modulate the synapses in neuronal networks and particularly on the synapses. To study synaptic stripping in a physiologically relevant system, we used microfluidic devices that allow to reconstitute *in vitro* mature cortico-cortical networks [34–36]. The microfluidic device consists of a presynaptic and a postsynaptic compartment containing both cortical neurons. An intermediate synaptic compartment receives axons from pre-synaptic cortical neurons and dendrites originating from post-synaptic cortical neurons. The three compartments are connected by 3-µm-wide microchannels that are 500 µm long for axons and 75 µm long for dendrites **(Fig. 5C)**. Because the axonal channels are 500 µm long, only axons from the cortex can reach the synaptic compartment. A laminin gradient from the pre-synaptic chamber to the post-synaptic chamber limits the number of post-synaptic axons that can reach the synaptic chamber **(Fig. 5C)**. Microglial cells were labeled with a GFP-encoding lentivirus and introduced in the synaptic compartment of the microfluidic chamber. We assessed the co-culture by staining all the compartments with antibodies recognizing GFP, a dendritic microtubule-associated protein 2 (MAP2), and an axonal phosphorylated neurofilament H (SMI31) marker **(Fig. 5D)**. At DIV10, previous reports have shown that the circuit achieves functional maturity, as defined by kinetics of neurite outgrowth, synapse formation and function, axonal transport, and neural activity [35,36]. At this stage, pre-synaptic cortical neurons have established functional excitatory connections to post-synaptic cortical neurons and pre-treated GFP-microglial cells were introduced into the synaptic compartments. All microfluidic experiments were performed from DIV10 to DIV11 for cortico-cortical neuronal networks and 24 hours after chemical and mechanical treatments of GFP-microglial cells.

We stained synaptic compartments with antibodies recognizing pre-synaptic protein synaptophysin (SYN) and the postsynaptic density protein 95 (PSD95). Confocal Z-stacks images of 3 µm depth representing the entire layer of synaptic connections into the synaptic compartment were acquired at different zones and acquisitions of different areas containing at least one GFP-microglia and pre- and post-synaptic proteins labelled were used for quantifications (**Supplementary Movie S6**). Images were first thresholded to remove non-specific signals and then the number of synaptophysin and PSD-95 spots were counted automatically by using the Analyze Particle plugin in FIJI [37]. Inside and outside areas were delimited by masks of the contour of microglial cells **(Fig. 5E-F)**. Our findings showed that the ratio between inside and outside post-synaptic dot density (PSD-95) was significantly lower in mechanically-activated microglial cells (0.5±0.2) than in LPS-treated (1.0±0.2) and control (0.8±0.3) cells **(Fig. 5G)**. Interestingly, we observed similar results for the ratio between inside and outside pre-synaptic dot density (synaptophysin) that was significantly decreased in mechanically-activated microglial cells (0.9±0.3) than in LPS-treated (1.3±0.2) and control (1.4±0.4) cells. Altogether, these results suggest that stretch injured-microglial cells have an enhanced stripping activity.

To confirm these results, we performed confocal z-stack images in high-resolution mode of microglial cells (**Supplementary Movie S7**). Then, we subtracted a basal focal plane of 3 µm thick, corresponding to the mean thickness of the synapses resting in the synaptic compartment **(Fig. 5I and Supplementary Movie S8)** to observe synaptic proteins only localized within the microglial cells **(Fig. 5J)**. Then, we compared the ratio between inside and outside synaptophysin and PSD95 dot density. We found a higher PSD95 protein dot ratio in mechanically-activated microglial cells (11.2±9.3) than in LPS-activated (2.7±0.6) and control (3.4±1.5) cells **(Fig. 5K-L)**. In addition, our results showed a higher synaptophysin protein dot ratio in mechanically-activated microglial cells (11.9±8.1) than in LPS-activated (2.6±3) and healthy cells (3.1±1.5).

Altogether, these results demonstrated that, unlike LPS treatment, a single mechanical stretch on microglial cells induces an enhanced synaptic stripping activity on healthy neuronal networks. Knowing that microglial cells can remain *in vivo* in an activated state for very long periods, our findings are therefore very important to better understand the remodeling of the neuronal connectivity after a traumatic event.

## 3. Discussion

Brain injuries are complex and heterogeneous pathologies that implicates many protagonists. Microglial cells are rapidly activated after a brain injury and many released signals collectively trigger an inflammatory cascade in brain tissues and peripheral immune cells can be recruited within minutes following injury [38]. We demonstrate here that microglial cells are mechanosensitive cells and change their morphology, cytoskeleton, and phagocytic activity in response to a stretching deformation, as observed during brain injury events.

A short and moderate stretch deformation leads to a reactive state, which is characterized by higher Iba1 levels. LPS-activated microglial cells secrete a wide range of pro-inflammatory cytokines that were not secreted by stretched cells. Our findings indicated that a stretch injury does induce the secretion of pro-inflammatory cytokines, but only the concentration of TNF-α, which has been shown to influence synaptic scaling, postsynaptic current frequency, and synaptic plasticity [39–41]. It is therefore interesting to note that a notable difference exists in the activation state of microglia in response to chemical or mechanical treatments, suggesting that different pathways are used in both situations.

In addition to important differences in the secretory profile, stretch insults induce abundant double-strand DNA breaks, whereas DNA integrity was not impacted by LPS-treatment. Interestingly, the confined migration of microglial cells in narrow spaces does not induce DNA damages, suggesting that microglia are able to discriminate between endogenously induced mechanical stress during migration and exogenously induced stress during short and mild injury.

Phagocytosis is part of the innate immune response of microglia, but also mediates the adaptive responses by contributing to antigen presentation [42]. A mechanical injury enhances phagocytosis in microglial cells, which is often considered as beneficial for tissue homeostasis by rapidly clearing dying cells, preventing the spillover of proinflammatory and neurotoxic molecules [43]. However, different targets and related receptors can finely tune microglia responses, which appear as a continuum of activation states [44]. For instance, phagocytosis of apoptotic neurons mediated by microglial triggering receptor expressed on myeloid cells-2 (TREM-2) was associated with decreased production of pro-inflammatory cytokines [45], while myelin debris phagocytosis enhanced the pro-inflammatory and dampened the anti-inflammatory profile in microglia [46].

We provide evidence that mechanically-activated microglia can remove damaged cells as well as stripping synapses from neurons. Activation of microglia in response to a stretch injury can protect neurons by removal of inhibitory synapses. Further studies will be required to develop methods that distinguish the phagocytic role of activated microglia by removing dying cells from their neuroprotective role by stripping synapses. Separating the neuroprotective and phagocytic phenotypes of activated microglia, will allow to decipher the molecular signature for protective microglia. It will be also interesting to understand the duration of the neuroprotection provided by mechanically-activated microglia, as well as its reversibility, by studying for instance if neurons can instruct activated microglia to transition from a neuroprotective to a phagocytic phenotype.

## 4. Conclusion

Microglial are sensitive cells that react and adapt to chemical and mechanical treatments in different ways. The mechanical activation of microglial cells must be considered as a key stage in neuroinflammation and synaptic stripping in the hours and days following the lesion. Our results define a new role for mechanically-activated microglia beyond being a mere biochemically alerted pathologic sensor. Collectively, these data raise the possibility that the activation of microglia in injured brains leads to an enhanced protective role in the injured brain. It will be interesting in future works to investigate how changes in chromatin compaction and DNA damage in stretched microglia could lead to genomic instabilities. Despite the fact that the integrin (α5β1)/FAK pathway has been recently recognized as an important contributing pathway in stretched microglial responses[47], the mechanotransduction mechanism by which stretch induces morphological changes, chromatin compaction and enhanced stripping activity is not well understood. Further analyses of the molecular pathways involved in the mechanical activation process are therefore required to better identify mechanotransduction pathways in stretched microglia and to develop new therapeutic strategies for preventing long-term disabilities after brain trauma.

## 5. Experimental Section

### Preparation of the cell culture substrate

To reproduce mechanical deformations observed during TBI events, stretchable chambers were made with a thin polydimethylsiloxane (PDMS) which has increasingly been employed for the fabrication of neuronal cell culture platforms and microfluidic [48,49]. The PDMS curing agent (Dow Corning, Sylgard 184) was mixed with a base agent in a mass ratio of 12:1 in 15 mL centrifugal tubes. The mixture was placed in a vacuum for 30 min to remove air bubbles and was then transferred on the top of a silanized Teflon mold. Then, the PDMS mixture was spin-coated (POLOS Wafer Spinner) with a speed gradually increasing from 100 to 600 rpm, for a total duration of 30 seconds and cured at 60 °C for 4 h. The final PDMS layer was 150 µm thick. The PDMS membrane was then stuck to a PDMS block (stretchable chamber). The field deformation of the elastic PDMS membranes was estimated by printing fluorescent protein (FITC-BSA) circles of 2000 µm2 on the membrane of the device. The device was then submitted to an uniaxial 20% stretch along the horizontal axis and the distances between the centers of the circles were determined along horizontal and vertical axes [23].

### Microfluidic device fabrication

The design of polydimethylsiloxane microfluidic devices is well documented [34–36]. Briefly, a master mold was made with SU-8 photoresist on silicon wafer using a dual thicknesses photolithography process. Indeed, the microfluidic circuit is composed of a set of thin and slender microchannels that connects three thicker culture chambers, namely *pre-synaptic*, *post-synaptic* and synaptic as shown in [36]. Epoxy resins (master replica of the 3 inches processed silicon wafers) were filled with the PDMS mixture. Air bubbles were removed by incubation in a desiccator under vacuum for 1 h, and polymerization was performed by incubating the PDMS for 3 h at 60°C. Finalized PDMS microchambers were cut and washed with 100% ethanol following by a quick passage through an ultrasonic bath and then washed with distilled water. Cut PDMS and glass-bottom, 0.17 µm-thick, 35 mm-diameter Petri dishes (FluoroDish, WPI) were placed into plasma cleaner under vacuum for 30 s for surface activation. After a rapid passage in an oven at 60 °C, PDMS pieces and Petri dishes were brought together to form an irreversible tight seal. The microfluidic devices were coated with a mixture of poly-*D*-lysine (0.1 mg/ml) in the upper and synaptic chambers, and with a mix of poly-*D*-lysine (0.1 mg/ml) + laminin (10 µg/ml) in the lower chamber overnight at 4 °C. Microchambers of microfluidic channels were washed 3 times with growing medium (Neurobasal medium supplemented with 2% B27, 2 mM Glutamax, and 1% penicillin/streptomycin) and placed at 37°C before neurons were plated.

### Cell culture and chemical/mechanical treatments

Microglial cells from the BV2 cell line (BV2, Elabscience, EP-CL-0493) were maintained in polystyrene T75 flasks in a cell culture incubator at 37 °C and 5% CO_2_. The BV2 cells were cultured in a proliferation medium composed of Dulbecco’s modified Eagle’s medium, high glucose (4.5 g/L) with L-glutamine (BE12-604F, Lonza) supplemented with 10% (v/v) fetal bovine serum (FBS; AE Scientific), and 1% penicillin and streptomycin antibiotics (AE Scientific). For all experimental groups, BV2 cells were seeded on stretched chamber of 10^3^ cells per chamber. All treatments (LPS and stretch) were performed 24 h after seeding and all experiments were done 24 h after treatment (48 h after seeding). Chemical-activation of microglial cells were done using Lipopolysaccharides from *Escherichia coli* (LPS) at 100 ng/ml concentration (Sigma-Aldrich, L4516-1MG). A single mechanical injury consisting of a 20% stretch was performed to the deformations occurring during traumatic brain injury (TBI). Stretching experiments was performed using an automatic stretcher (StrexCell STB-150) (**Fig. 3A**). To avoid the twisting/stretching of the chambers during their manipulation, a stabilizer was custom-made by 3D printing (Ulti Maker V1.9) that reinforced the structure of the stretchable chamber. It also permits to avoid any undesired deformations during the manipulation of the deformable chambers.

### Isolation and culture of primary microglial cells

Cortical microglia from CX3CR1^eGFP/+^ WT mice were isolated following experimental procedures described elsewhere [50]. In brief, brains of post-natal day 21 (P21) mice were dissected, and the midbrain, cerebellum and meninges were carefully removed. The remained tissue were placed in Dulbecco’s Modified Eagle’s Medium (DMEM, Sigma-Aldrich, Overijse, Belgium) supplemented with 1% penicillin/streptomycin (P/S, Invitrogen, Merelbeke, Belgium), followed by incubation with papain (17 U/mg, Sigma-Aldrich) and DNase I (10 mg/ml, Roche, Brussel, Belgium) for 30 min at 37 °C. Cell suspensions were filtered through a 70 µm cell strainer, centrifuged (5 min, 500 g) and pellets were resuspended in DMEM containing 30% stock isotonic Percoll (SIP, GE Healthcare, Diegem, Belgium). Hereafter, a density gradient was created by the addition of 70% SIP diluted in PBS and the suspension was centrifuged for 25 min at 650 g (brake 0, acceleration 4). The cell cloud at the interphase between 30% and 70% was collected, diluted in 10 ml cold PBS and centrifuged for 10 min at 500 g. Cell pellets were resuspended in magnetic activated cell sorting (MACS) buffer (2 mM EDTA and 0.5% fetal calf serum (FCS)) and microglia were isolated by positive selection using CD11b microbeads (Miltenyi Biotec, Gladbach, Germany), following the manufacturer’s instructions. CD11b^+^ cells were resuspended in DMEM supplemented with 10% FCS, 10% horse serum (Thermofisher, Waltham, MA, US) and 1% P/S (DMEM 10:10:1) and seeded onto stretchable chambers (30 x 10^3^ cells/well) pre-coated with poly-*D*-lysine (PDL, 20 µg/ml, Gibco, Waltham, MA, US) and collagen type IV (2 µg/ml, Sigma - Aldrich), and incubated in a humidified incubator at 37 °C and 5% CO_2_ for 7 days. Afterwards, a dynamic ramified morphology was induced by the addition of serum-free medium (hereafter referred as TIC medium) containing 5 µg/ml insulin, 5 µg/ml *N*-acetyl-cysteine, 100 µg/ml apo-transferrin, 0.1 µg/ml Na_2_SeO_3_, 1 µg/ml heparan sulfate, 2 µg/ml human TGF-ß (PeproTech, Rocky Hill, NJ, US), 0.1 µg/ml murine IL-34 (BioL egend, Amsterdam, The Netherlands), 1.5 µg/ml ovine wool cholesterol, 3 µg/ml *L*-glutamine in DMEM/F12. For all experiments, cells were seeded 7 days in DMEM 10:10:1 medium followed by 3-7 days TIC medium before experiments.

### Primary rat cortical neurons cultures

Primary cortical neurons were prepared as previously described.[51] Briefly, cortex was dissected from E15.5 wild-type (Wistar-Han) rat embryos, then digested with a papain and cysteine solution followed by two incubations with trypsin inhibitor solutions, and finally dissociated mechanically. Dissociated cortical neurons were resuspended in growing medium (5×10^6^ cells in 80 µl) and plated in the chamber with a final density of ∼7000 cells/mm^2^. Cortical neurons were plated first on the upper chamber followed by addition of growing medium in the synaptic chamber. Striatal neurons were then added in the lower chamber. Neurons were place in the incubator for at least 3 hours, and all compartments were gently filled with growing medium. Microchambers are then carefully inspected to avoid any cell contamination in the synaptic chamber before experiments.

### Lentiviruses

BV2 microglial cells were infected with lentiviruses (LV) before to be plated in stretchable chambers for 24 h. The following LV construct was used for the study: Lenti pSIN GFP (Gene ID: 7011691; Type A).

### Indentation and measurements protocol

Microglial cells stiffness was tested using a Chiaro indenter system (Optics11, Amsterdam, the Netherlands). It consists of a ferrule-top force transducer[52] composed of a micromachined cantilever spring with an optical fiber readout, was mounted on a 3D printed holder screwed to a Z-piezoelectric actuator (PI p-603.5S2, Physik Instrumente). The single-mode fiber of the readout was coupled to an interferometer (OP1550, Optics11), where the interference signal was directly translated into cantilever deflection. The piezoelectric actuator with the probe was mounted on a XYZ micromanipulator (PatchStar, Scientifica) for automatic mapping of mechanical properties. Indentation mapping was performed in parallel lines, with at least 25 (5X x 5Y) points per chamber. Distance between two adjacent locations were 15 µm, to ensure that two indentations areas were sufficiently far away from the other. Colloidal probes with a tip diameter of 3 µm were used for testing the microglial cells stiffness. Three samples were tested for each condition. A total of 110-126 indentation data points were recorded per experimental group. Before testing, the sensitivity calibration of the cantilever was conducted by indenting a hard glass surface. The Hertz model was used to fit an initial loading data up to the cantilever threshold value to obtain the true surface position:

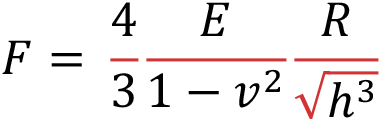

where F is the load, E is an elastic modulus, v is the Poisson’s ratio of compressibility (we assume that brain cells are incompressible v = 0.5), h is the indentation depth.

### Immunostaining in stretchable and microfluidic chambers

BV2 cells were fixed and permeabilized with 4% paraformaldehyde (Electron Microscopy Sciences) and 0.05% Triton X-100 (Sigma) in PBS (1X, Capricorn Scientific) for 15 min at room temperature (RT). The fixed cells were rinsed three times in warm PBS and incubated for 30 min with a blocking solution containing 1% BSA (GE Healthcare) and 5% FBS in PBS. BV2 cells were labelled for F-actin with Alexa Fluor 555 phalloidin (1:200; Invitrogen A34055), DNA with DAPI (1:200; Thermo Fisher Scientific, D1306), and Iba1 (1:750) Sopachem, 019-19741). γH2Ax was labelled with Anti-phospho Histone H2A.X (Ser139), coupled to Alexa Fluor 488 Conjugate Antibody monoclonal antibody (1:200; Millipore Sigma, clone JBW301: 05-636-AF488) for 1 hour at RT. PDMS membranes with immunostained cells were cut off the stretch chamber and mounted on microscope slides with slow-fade diamond antifade (Thermo Fisher, Molecular Probes) for epifluorescence and confocal imaging. Neurons in the microchambers were fixed with a PFA/Sucrose solution (4%/4% in PBS) for 20 min at RT. The fixation buffer was rinsed three times with PBS and neurons were incubated for 1h at RT with a blocking solution (BSA 1%, normal goat serum 2%, Triton X-100 0.1%). The compartment of interest was then incubated with primary antibodies overnight at 4 °C and appropriate fluorescent secondary antibodies were incubated for 1 h at RT. The immunofluorescence was maintained in PBS for a maximum of one week in the dark at 4 °C. The following primary antibodies were used: PSD95 (Millipore, MAB1598, 1:1,000), Synaptophysin (Abcam, AB14692, 1:200), MAP2 (Chemicon, AB5622, 1:500), GFP (Abcam, Ab13970, 1:2,000).

### Cytokine quantification

Meso Scale Discovery (MSD) electrochemiluminescence multiplex immunoassay (Meso Scale Diagnostics, Maryland, US) was performed to quantify the amount of inflammatory cytokines and chemokines released by control, LPS-treated and mechanically-activated BV2 microglial cells (V-PLEX Plus Proinflammatory Panel1 Mouse Kit, K15048G-1). The MSD kit permitted to quantify the following cytokines: interferon gamma (IFN-γ); interleukins 2, 4, 5, 6, 10, 12p70, and 1ß (IL-4, IL-5, IL-6, Il-10, and IL-1ß); tumour necrosis factor alpha (TNF-α) and the chemokine KC/GRO also known as CXCL1, even at a very low concentration (lowest LLOD is 0.65 pg/ml for the IFN-γ). The multi-array technology combines electrochemiluminescence and multi-spot plates to enable precise quantitation of multiple analytes in a single sample requiring less time and effort than other assay platforms. The assay can be considered as a “sandwich immunoassay” with a 96-well 10 spot-plate pre-coated with capture antibodies.

### Migration assays

Microglial cells were chemically or mechanically treated in stretchable chambers and trypsinized 24 hours post-treatment. Cells were seeded on PDMS-coated fluorodish that were microprinted with mix of Poly-L-Lysine and Laminine (PLL-LA) microstripes of 15 µm wide. After minimum 4 hours of cell spreading, fluorodishes were placed under a microscope at 37°C - 5% CO_2_. Images were taken every 10 minutes for 15 hours and time-lapse sequences were analyzed with FIJI and the Cell Tracker code on MatLab to determine the migration speed and the persistence time.

### Phagocytosis assay

Microglial cells were chemically (LPS) or mechanically (stretch) treated in stretchable chambers. After 24 hours post-treatment, a controlled number (10 beads/cell) of fluorescent latex beads of 1 µm of diameter (Sigma-Aldrich, L2778-1ML) were introduced in the medium for 1 hour. The medium was then removed to eliminate non-phagocytosed beads and cells were fixed with PFA for 15 minutes. After immunostaining, Images were taken with confocal microscope on, at least, 3 different regions of interest (ROIs) per sample.

### Synaptophysin and PSD95 analysis

Colocalization and independent dots analyses of synaptophysin and PSD95 were performed using ImageJ. Airyscan images were thresholded to remove non-specific signals. The number of synaptophysin spots overlapping, juxtaposed, or separated by no more than 2 pixels (130 nm) to PSD95 spots were counted automatically. Results were expressed as a function of density outside the area of microglial cells and the density inside. Each condition was tested using at least 3 chambers per culture from 3 independent cultures. In each chamber, 3 fields were analyzed in which 3 regions of interest were selected (n = number of fields).

### Epifluorescence, confocal and time-lapse imaging

Immunostained preparations of BV2 cells were observed in epifluorescence and confocal modes with a Nikon A1R HD25 (Nikon, Japan) motorized inverted microscope equipped with ×20, ×40 and ×60 Plan Apo (numerical aperture, 1.45; oil immersion) objectives and lasers that span the violet (405 and 440 nm), blue (457, 477 and 488 nm), green (514 and 543 nm), yellow-orange (568 and 594 nm) and red (633 and 647 nm) spectral regions. Epifluorescence images were recorded with a Prime 95B camera (Photometrics) using NIS-Elements Advanced Research 4.5 software (Nikon). Z-stack images were collected using ×60 or ×40 objective for three channels (DAPI, TRITC and FITC) from the entire volume of the nuclei using a step size of 0.15 µm or the cell using a step size of 1 µm. For the cell exposure times and the laser power were kept constant, and the acquired stack of images was deconvolved to remove the focus light. PSD95/SYN immunostaining were acquired in the synaptic chamber with a ×63 oil-immersion objective (1.4 NA) using an inverted confocal microscope (LSM 710, Zeiss) coupled to an Airyscan detector to improve signal-to-noise ratio and to increase spatial resolution. Time-lapse experiments were carried out at 37°C and 5% CO2 for 15 hours on a Nikon A1R HD25 (Nikon, Japan) motorized inverted microscope equipped with a cage incubator (OkoLab) and controlled with the NIS Elements Advanced Research 4.0 software (Nikon, Japan).

### Image analysis

All images were acquired with NIS-Elements Advanced Research software (v.4.5, Nikon, Japan) using similar illumination and recording conditions (camera frequency, gain and lamp intensity). The quantification of Iba1 and actin fluorescence intensity was performed using a corrected total fluorescence intensities analysis method. The area, the raw integrated density, the mean grey value, and the number of cells were measured for each image. In addition, five random background regions were selected to obtain a mean grey value of the fluorescent background[53]. The corrected total fluorescence was calculated using the following equation:

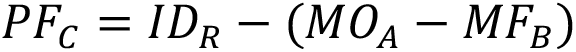

with PF_C_ was the corrected protein fluorescence intensity, ID_R_ the raw integrated density, MO_A_ the marked objects area and MF_B_ the mean fluorescence background. Images were taken at least at 3 different regions of interest (ROI) for each sample obtained from 6 different chambers (n=ROIs). Confocal images were processed using FIJI software[37].

### Statistical analysis

Experimental data were presented using a boxplot, multiple-variable or histogram representation and statistical comparisons were realized by either a Student t-test or ANOVA, with ns ≥ 0.05, 0.01 ≤ p* ≤ 0.05, 0.001 ≤ p** ≤ 0.01, p*** ≤ 0.001.

### Ethical compliance

Experiments in mice were conducted in accordance with the European Community guiding principles on the care and use of animals and with the approval of the Ethical Committee on Animal Research of Hasselt University (Project licence 201956K). Animals were group-housed in a temperature and humidity-controlled room with ad libitum access to food and water and a 12 h light–dark cycle. CX3CR1^eGFP/+^ were obtained by breeding CX3CR1^eGFP/GFP^ with wild type C57BL6 mice. CX3CR1^eGFP/GFP^ mice[54] were acquired from the European Mouse Mutant Archive (EMMA) Institute with the approval of Steffen Jung (Weizmann Institute of Science).

## Supporting Information

Supporting information is available from the Wiley Online or from the author

## Acknowledgements

The authors thank Djamal Achour from the IMPact de l’Environnement Chimique sur la Santé (ULR 4483 - IMPECS) team for his help with the quantification of cytokines and Johanna Cormenier for her help with microfluidic experiments. A.P. and S.G. acknowledges funding from FEDER Prostem Research Project no. 1510614 (Wallonia DG06), the F.R.S.-FNRS Epiforce Project no. T.0092.21, the F.R.S.-FNRS CellSqueezer Project no. J.0061.23, the F.R.S.-FNRS Optopattern Project no. U.NO26.22 and the Interreg MAT(T)ISSE project, which is financially supported by Interreg France-Wallonie-Vlaanderen (Fonds Européen de Développement Régional, FEDER-ERDF). Y.A.A was supported by an FWO senior postdoctoral fellowship (12H8220N). Y.A.A. and BB are supported by the BOF small research project 21KP06BOF. F.S. acknowledges funding from European Research Council (ERC) under the European Union’s Horizon 2020 research and innovation programme AdG grant agreement no. 834317, Fueling Tranport. We acknowledge Yasmina Saoudi and the Photonic Imaging Center of Grenoble Institute Neuroscience (Univ Grenoble Alpes – Inserm U1216) which is part of the ISdV core facility and certified by the IBiSA label. A.P. is financially supported by F.R.S.-FNRS with a FRIA PhD grant and Fonds pour la Recherche Médicale en Hainaut (FRMH).

## Author contributions

S.G. and A.P. conceived the project. L.R. and S.G. supervised the project. A.P. performed microglial cell culture, stretching experiments, cell tracking, immunostaining and imaging. Y.A. and B.B. contributed to experiments using primary microglial cells. S.H. contributed to the quantification of cytokines. Experiments using microfluidic chambers were performed under the supervision of F.S. The article was written by S.G. and A.P, read and corrected by all authors, who all contributed to the interpretation of the results.

## Data Availability Statement

The data that support the findings of this study are available in the supplementary material of this article.

---

**Supplementary Figure 1.**
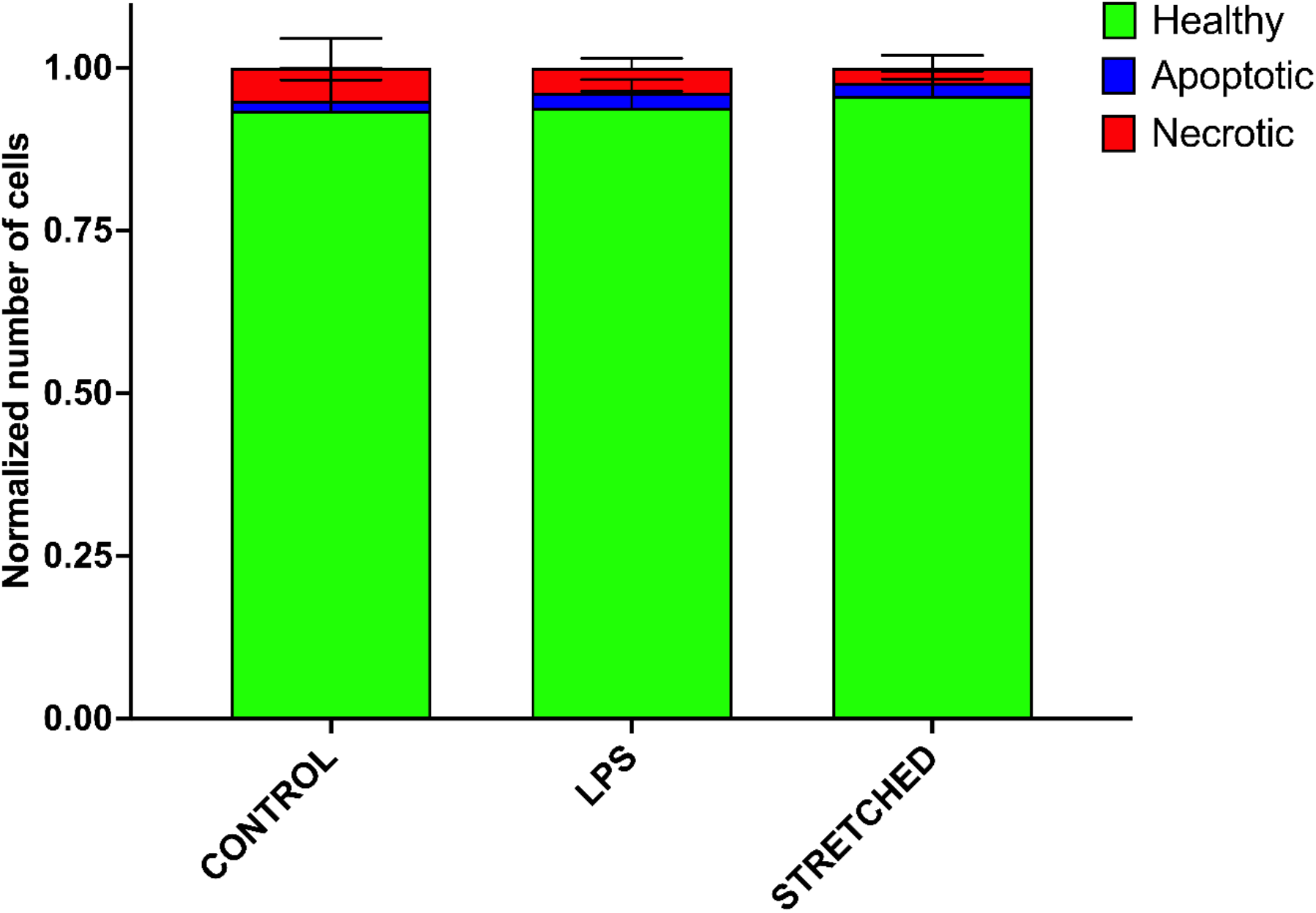
– Viability of microglial cells is not affected by chemical (LPS) and mechanical (stretch) treatments. The viability of BV2 cells was measured using Hoechst to label the DNA propidium iodide to label necrotic cells and Caspase 3/7 Green to label apoptotic cells with active caspase 3. Viability assays were performed 24 hours after treatments on fixed cells using 3 ROIs per sample, N=3 replicates (n.s. for each condition). The average total number of cells per ROI was 24 cells for Control, 74 cells for LPS and 63 cells for Stretched.

**Supplementary Figure 2.**
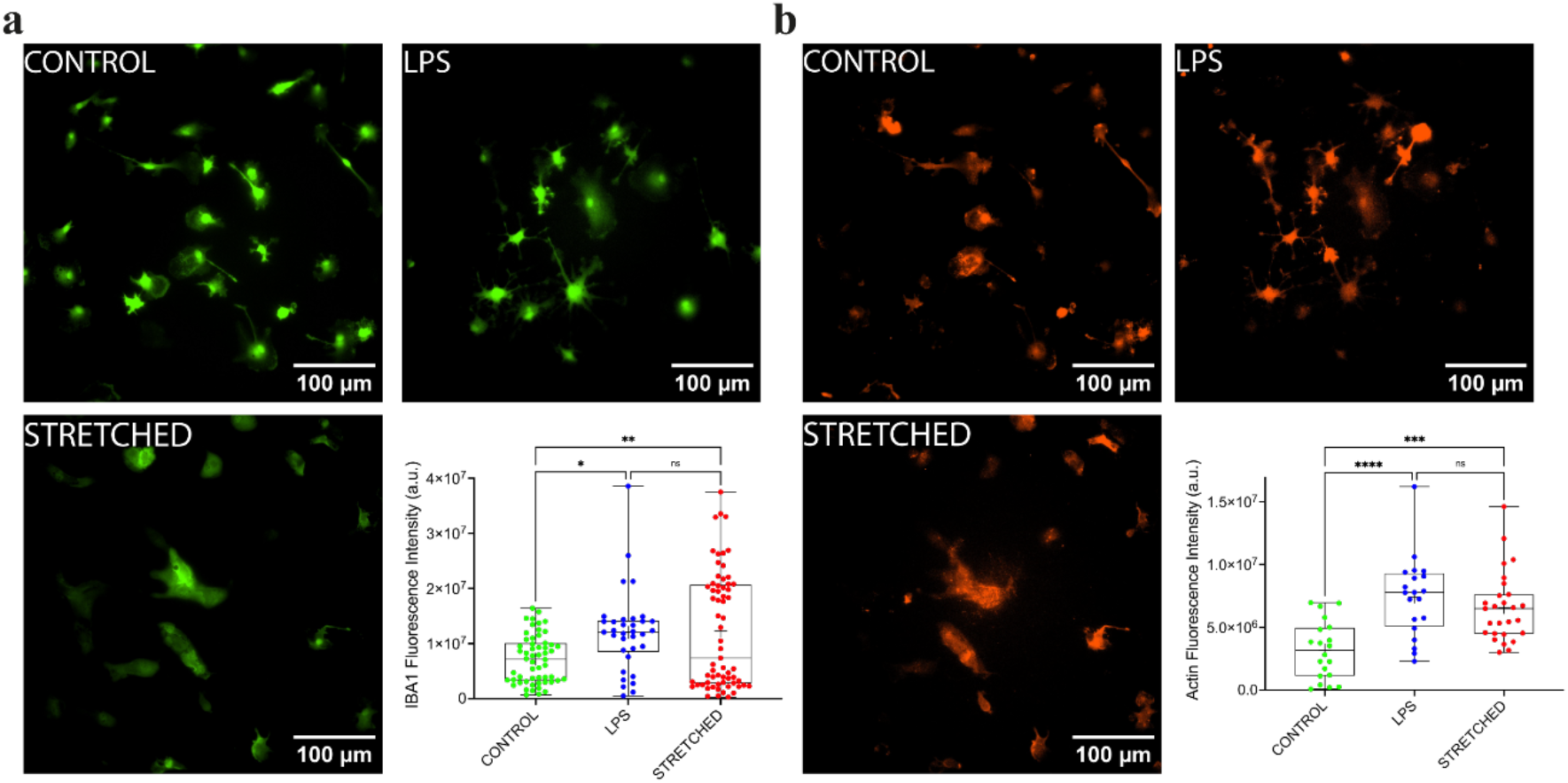
– Chemical and mechanical activation of primary mouse microglial cells *in vitro*. (A) Fluorescent images of immunostained primary microglial cells (IBA1 in green) with no treatment (control, in green), LPS-treatment at 100 ng/ml for 24 hours (LPS, in blue) and mechanical activation with a 20% stretch (stretched, in red). Iba1 fluorescence intensity for control (n=56, in green), LPS-treated (n=34, in blue) and stretched (n=65, in red) cells. (B) Fluorescent images of immunostained primary microglial cells (F-actin in red) for control, LPS-treated and stretched cells. Actin fluorescence intensity for control (n=33, in green), LPS-treated cells (n=70, in blue) and stretched (n=59, in red) cells. Experiments for control and LPS were performed on 5 chambers and on 6 chambers for stretched cells using 3 different cultures. Scale bars are 100 μm. ns is not significant, 0.01 ≤ p^∗^ ≤ 0.05, 0.001 ≤ p^∗∗^ ≤ 0.01, p^∗∗∗^ ≤ 0.001.

**Supplementary Figure 3.**
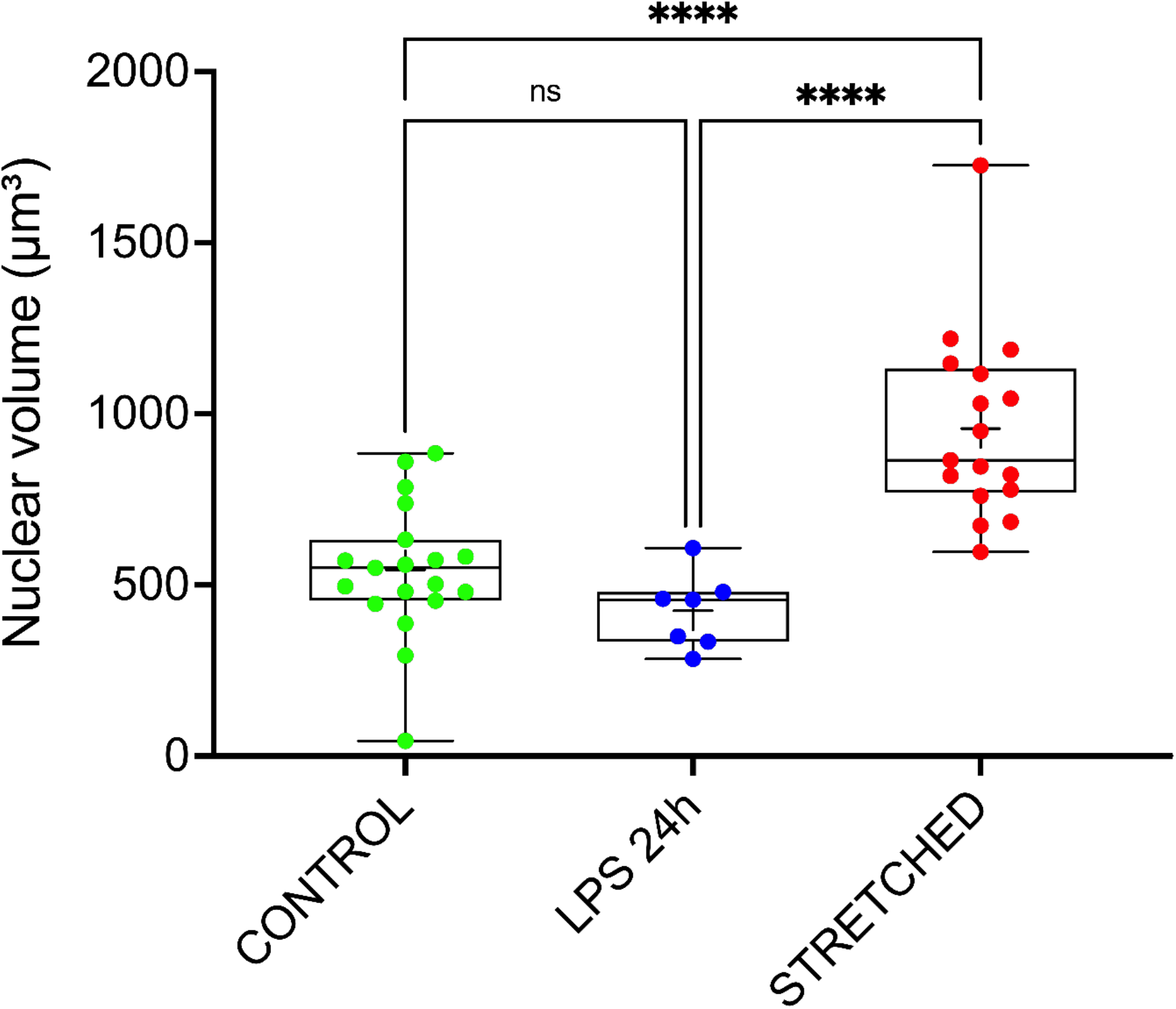
– The nuclear volume increased in stretched microglia. Nuclear volume for control (n=19, in green), LPS-treated (n=7, in blue) and stretched (n=17, in green) cells (N≥3 replicates). ns is not significant, 0.01 ≤ p^∗^ ≤ 0.05, 0.001 ≤ p^∗∗^ ≤ 0.01, p^∗∗∗^ ≤ 0.001.

**Supplementary Figure 4.**
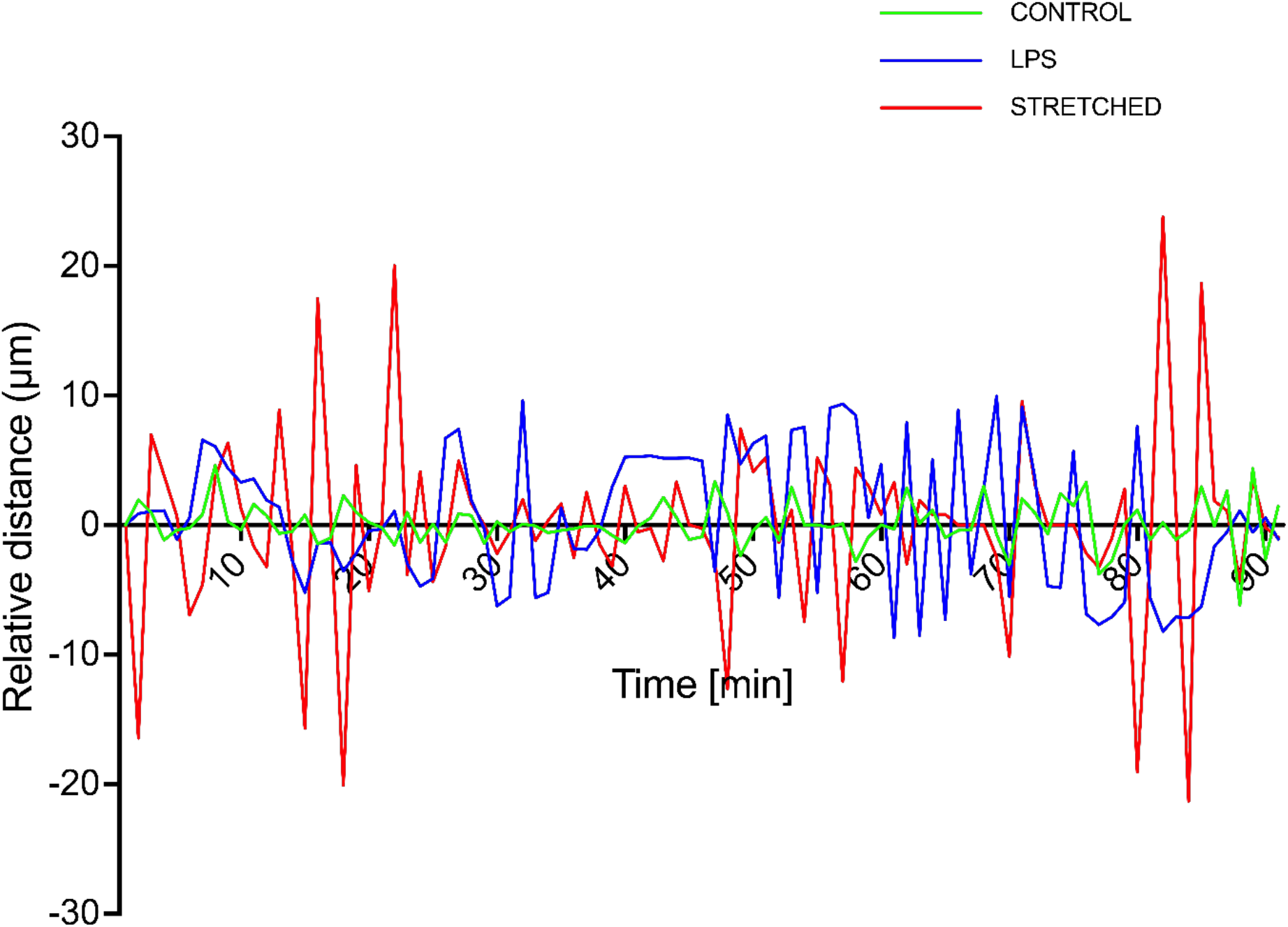
– The migration of stretched microglial cells is more persistent. Evolution of the travelling distance over time of control (in green), LPS-treated (in blue) and stretched (in red) microglial cells. Mechanical activation of microglial cells results in fewer directional changes and longer directional movements.

**Supplementary Movie S1 – Uniaxial stretching deformation.** An elastic PDMS chamber was fixed on a stretcher device placed in a biological safety cabinet to ensure sterility. The chamber was stretched from the rest position to the 20% stretch position in less than a second, then maintained at the 20% stretch position for 1 second and relaxed to the rest position in less than a second. Stretched cultures were then placed at 37°C and 5% CO_2_ for 24 hours before any further experiments.

**Supplementary Movie S2 – Determination of the mechanical stiffness of microglial cells.** The elastic modulus of microglial cells was probed inside the stretchable chamber using a nanoindenter equipped with an automatic cell surface detection mode. The probe navigated 15 µm along x- and y-axis to avoid repetitive measurement on the same cell.

**Supplementary Movie S3 – Migration of microglial cells on proteins microstripes.** Time-lapse movie in DIC mode of BV2 microglial cells migrating on adhesive microstripes. The duration time is 15 hours.

**Supplementary Movie S4 – Migration of microglial cells on proteins microstripes.** Time-lapse movie in DIC mode of BV2 microglial cells migrating on adhesive microstripes. Nuclei are stained with Hoechst (blue) and microstripes with PLL-FITC (green).

**Supplementary Movie S5. 3D confocal view of the nucleus in a stretched microglial cell.** Nuclei were stained with DAPI (in blue) and DNA damages (in red) resulting from the stretch injury were immunolabelled with γH2Ax.

**Supplementary Movie S6 – Confocal view of basal plane where stretched microglial cells interact with synapses.** Z-projection of a 3 µm thick confocal section (basal plane) containing the synapses resting in the synaptic compartment but not those phagocyted by a stretched microglial cell. Synaptophysin is in red, PSD95 in purple and GFP in green.

**Supplementary Movie S7 – Confocal view of the presence of synapses inside and outside stretched microglial cells.** Z-projection of a 15 µm thick confocal image containing the synapses resting in the synaptic compartment and those phagocyted by a stretched microglial cell. Synaptophysin is in red, PSD95 in purple and GFP in green.

**Supplementary Movie S8 – Confocal view of the presence of synapses within a stretched microglial cell.** Z-projection of a confocal image from which a basal section of 3 µm thick (see Supplementary Movie S6) was substracted. The projection shows only the synapses phagocyted by a stretched microglial cell. Synaptophysin is in red, PSD95 in purple and GFP in green.

## Bibliography

1. Tyler, W.J. (2012) The mechanobiology of brain function. Nat Rev Neurosci, 13 (12), 867–878.

2. Gupta, R., and Sen, N. (2016) Traumatic brain injury: a risk factor for neurodegenerative diseases. Reviews in the Neurosciences, 27 (1), 93–100.

3. Goddery, E.N., Fain, C.E., Lipovsky, C.G., Ayasoufi, K., Yokanovich, L.T., Malo, C.S., Khadka, R.H., Tritz, Z.P., Jin, F., Hansen, M.J., and Johnson, A.J. (2021) Microglia and Perivascular Macrophages Act as Antigen Presenting Cells to Promote CD8 T Cell Infiltration of the Brain. Front. Immunol., 12, 726421.

4. Janda, E., Boi, L., and Carta, A.R. (2018) Microglial Phagocytosis and Its Regulation: A Therapeutic Target in Parkinson’s Disease? Front. Mol. Neurosci., 11, 144.

5. Marín-Teva, J.L., Dusart, I., Colin, C., Gervais, A., van Rooijen, N., and Mallat, M. (2004) Microglia Promote the Death of Developing Purkinje Cells. Neuron, 41 (4), 535–547.

6. Halder, S.K., and Milner, R. (2019) A critical role for microglia in maintaining vascular integrity in the hypoxic spinal cord. Proc. Natl. Acad. Sci. U.S.A., 116 (51), 26029– 26037.

7. Paolicelli, R.C., Bolasco, G., Pagani, F., Maggi, L., Scianni, M., Panzanelli, P., Giustetto, M., Ferreira, T.A., Guiducci, E., Dumas, L., Ragozzino, D., and Gross, C.T. (2011) Synaptic Pruning by Microglia Is Necessary for Normal Brain Development. Science, 333 (6048), 1456–1458.

8. Smolders, S.M.-T., Kessels, S., Vangansewinkel, T., Rigo, J.-M., Legendre, P., and Brône, B. (2019) Microglia: Brain cells on the move. Progress in Neurobiology, 178, 101612.

9. Jurga, A.M., Paleczna, M., and Kuter, K.Z. (2020) Overview of General and Discriminating Markers of Differential Microglia Phenotypes. Front. Cell. Neurosci., 14, 198.

10. Loane, D.J., Kumar, A., Stoica, B.A., Cabatbat, R., and Faden, A.I. (2014) Progressive Neurodegeneration After Experimental Brain Trauma: Association With Chronic Microglial Activation. Journal of Neuropathology & Experimental Neurology, 73 (1), 14–29.

11. Smith, C. (2013) Review: The long-term consequences of microglial activation following acute traumatic brain injury: Neuroinflammation after trauma. Neuropathology and Applied Neurobiology, 39 (1), 35–44.

12. Lu, Y.-B., Franze, K., Seifert, G., Steinhauser, C., Kirchhoff, F., Wolburg, H., Guck, J., Janmey, P., Wei, E.-Q., Kas, J., and Reichenbach, A. (2006) Viscoelastic properties of individual glial cells and neurons in the CNS. Proceedings of the National Academy of Sciences, 103 (47), 17759–17764.

13. Bollmann, L., Koser, D.E., Shahapure, R., Gautier, H.O.B., Holzapfel, G.A., Scarcelli, G., Gather, M.C., Ulbricht, E., and Franze, K. (2015) Microglia mechanics: immune activation alters traction forces and durotaxis. Front. Cell. Neurosci., 9.

14. Henn, A. (2009) The suitability of BV2 cells as alternative model system for primary microglia cultures or for animal experiments examining brain inflammation. ALTEX, 83–94.

15. Wang, Y., Jin, G., Miao, H., Li, J.Y.-S., Usami, S., and Chien, S. (2006) Integrins regulate VE-cadherin and catenins: Dependence of this regulation on Src, but not on Ras. Proceedings of the National Academy of Sciences, 103 (6), 1774–1779.

16. Hoogland, I.C.M., Houbolt, C., van Westerloo, D.J., van Gool, W.A., and van de Beek, D. (2015) Systemic inflammation and microglial activation: systematic review of animal experiments. J Neuroinflammation, 12 (1), 114.

17. Lund, S., Christensen, K.V., Hedtjärn, M., Mortensen, A.L., Hagberg, H., Falsig, J., Hasseldam, H., Schrattenholz, A., Pörzgen, P., and Leist, M. (2006) The dynamics of the LPS triggered inflammatory response of murine microglia under different culture and in vivo conditions. Journal of Neuroimmunology, 180 (1–2), 71–87.

18. Lively, S., and Schlichter, L.C. (2018) Microglia Responses to Pro-inflammatory Stimuli (LPS, IFNγ+TNFα) and Reprogramming by Resolving Cytokines (IL-4, IL-10). Front. Cell. Neurosci., 12, 215.

19. Sasaki, Y., Ohsawa, K., Kanazawa, H., Kohsaka, S., and Imai, Y. (2001) Iba1 Is an Actin-Cross-Linking Protein in Macrophages/Microglia. Biochemical and Biophysical Research Communications, 286 (2), 292–297.

20. Rheinlaender, J., Dimitracopoulos, A., Wallmeyer, B., Kronenberg, N.M., Chalut, K.J., Gather, M.C., Betz, T., Charras, G., and Franze, K. (2020) Cortical cell stiffness is independent of substrate mechanics. Nat. Mater., 19 (9), 1019–1025.

21. Beattie, E.C., Stellwagen, D., Morishita, W., Bresnahan, J.C., Ha, B.K., Von Zastrow, M., Beattie, M.S., and Malenka, R.C. (2002) Control of Synaptic Strength by Glial TNFα. Science, 295 (5563), 2282–2285.

22. Henning, L., Antony, H., Breuer, A., Müller, J., Seifert, G., Audinat, E., Singh, P., Brosseron, F., Heneka, M.T., Steinhäuser, C., and Bedner, P. (2023) Reactive microglia are the major source of tumor necrosis factor alpha and contribute to astrocyte dysfunction and acute seizures in experimental temporal lobe epilepsy. Glia, 71 (2), 168–186.

23. Lantoine, J., Procès, A., Villers, A., Halliez, S., Buée, L., Ris, L., and Gabriele, S. (2021) Inflammatory Molecules Released by Mechanically Injured Astrocytes Trigger Presynaptic Loss in Cortical Neuronal Networks. ACS Chem. Neurosci., 12 (20), 3885–3897.

24. Schaks, M., Giannone, G., and Rottner, K. (2019) Actin dynamics in cell migration. Essays in Biochemistry, 63 (5), 483–495.

25. Lammerding, J. (2011) Mechanics of the Nucleus, in Comprehensive Physiology, 1ed., Wiley, pp. 783–807.

26. Kalukula, Y., Stephens, A.D., Lammerding, J., and Gabriele, S. (2022) Mechanics and functional consequences of nuclear deformations. Nat Rev Mol Cell Biol.

27. Szczesny, S.E., and Mauck, R.L. (2017) The Nuclear Option: Evidence Implicating the Cell Nucleus in Mechanotransduction. Journal of Biomechanical Engineering, 139 (2).

28. Long, J.T., and Lammerding, J. (2021) Nuclear Deformation Lets Cells Gauge Their Physical Confinement. Developmental Cell, 56 (2), 156–158.

29. Luciano, M., Xue, S.-L., De Vos, W.H., Redondo-Morata, L., Surin, M., Lafont, F., Hannezo, E., and Gabriele, S. (2021) Cell monolayers sense curvature by exploiting active mechanics and nuclear mechanoadaptation. Nat. Phys., 17 (12), 1382–1390.

30. Versaevel, M., Grevesse, T., and Gabriele, S. (2012) Spatial coordination between cell and nuclear shape within micropatterned endothelial cells. Nat Commun, 3 (1), 671.

31. Kuo, L.J., and Yang, L.-X. (2008) Gamma-H2AX - a novel biomarker for DNA double-strand breaks. In Vivo, 22 (3), 305–309.

32. Hochreiter-Hufford, A., and Ravichandran, K.S. (2013) Clearing the Dead: Apoptotic Cell Sensing, Recognition, Engulfment, and Digestion. Cold Spring Harbor Perspectives in Biology, 5 (1), a008748–a008748.

33. Brown, G.C., and Neher, J.J. (2014) Microglial phagocytosis of live neurons. Nat Rev Neurosci, 15 (4), 209–216.

34. Scaramuzzino, C., Cuoc, E.C., Pla, P., Humbert, S., and Saudou, F. (2022) Calcineurin and huntingtin form a calcium-sensing machinery that directs neurotrophic signals to the nucleus. Sci. Adv., 8 (1), eabj8812.

35. Moutaux, E., Christaller, W., Scaramuzzino, C., Genoux, A., Charlot, B., Cazorla, M., and Saudou, F. (2018) Neuronal network maturation differently affects secretory vesicles and mitochondria transport in axons. Sci Rep, 8 (1), 13429.

36. Virlogeux, A., Moutaux, E., Christaller, W., Genoux, A., Bruyère, J., Fino, E., Charlot, B., Cazorla, M., and Saudou, F. (2018) Reconstituting Corticostriatal Network on-a-Chip Reveals the Contribution of the Presynaptic Compartment to Huntington’s Disease. Cell Reports, 22 (1), 110–122.

37. Schindelin, J., Arganda-Carreras, I., Frise, E., Kaynig, V., Longair, M., Pietzsch, T., Preibisch, S., Rueden, C., Saalfeld, S., Schmid, B., Tinevez, J.-Y., White, D.J., Hartenstein, V., Eliceiri, K., Tomancak, P., and Cardona, A. (2012) Fiji: an open-source platform for biological-image analysis. Nat Methods, 9 (7), 676–682.

38. Corps, K.N., Roth, T.L., and McGavern, D.B. (2015) Inflammation and Neuroprotection in Traumatic Brain Injury. JAMA Neurol, 72 (3), 355.

39. Pascual, O., Ben Achour, S., Rostaing, P., Triller, A., and Bessis, A. (2012) Microglia activation triggers astrocyte-mediated modulation of excitatory neurotransmission. Proc. Natl. Acad. Sci. U.S.A., 109 (4).

40. Nguyen, P.T., Dorman, L.C., Pan, S., Vainchtein, I.D., Han, R.T., Nakao-Inoue, H., Taloma, S.E., Barron, J.J., Molofsky, A.B., Kheirbek, M.A., and Molofsky, A.V. (2020) Microglial Remodeling of the Extracellular Matrix Promotes Synapse Plasticity. Cell, 182 (2), 388–403.e15.

41. Clark, A.K., Gruber-Schoffnegger, D., Drdla-Schutting, R., Gerhold, K.J., Malcangio, M., and Sandkuhler, J. (2015) Selective Activation of Microglia Facilitates Synaptic Strength. Journal of Neuroscience, 35 (11), 4552–4570.

42. Litman, G.W., Cannon, J.P., and Rast, J.P. (2005) New Insights into Alternative Mechanisms of Immune Receptor Diversification, in Advances in Immunology, vol. 87, Elsevier, pp. 209–236.

43. Wolf, S.A., Boddeke, H.W.G.M., and Kettenmann, H. (2017) Microglia in Physiology and Disease. Annu. Rev. Physiol., 79 (1), 619–643.

44. Hanisch, U.-K., and Kettenmann, H. (2007) Microglia: active sensor and versatile effector cells in the normal and pathologic brain. Nat Neurosci, 10 (11), 1387–1394.

45. Takahashi, K., Rochford, C.D.P., and Neumann, H. (2005) Clearance of apoptotic neurons without inflammation by microglial triggering receptor expressed on myeloid cells-2. Journal of Experimental Medicine, 201 (4), 647–657.

46. Siddiqui, T.A., Lively, S., and Schlichter, L.C. (2016) Complex molecular and functional outcomes of single versus sequential cytokine stimulation of rat microglia. J Neuroinflammation, 13 (1), 66.

47. Shaughness, M.C., Pierron, N., Smith, A.N., and Byrnes, K.R. (2023) The Integrin Pathway Partially Mediates Stretch-Induced Deficits in Primary Rat Microglia. Mol Neurobiol.

48. Grevesse, T., Dabiri, B.E., Parker, K.K., and Gabriele, S. (2015) Opposite rheological properties of neuronal microcompartments predict axonal vulnerability in brain injury. Scientific Reports, 5 (1).

49. Lantoine, J., Grevesse, T., Villers, A., Delhaye, G., Mestdagh, C., Versaevel, M., Mohammed, D., Bruyère, C., Alaimo, L., Lacour, S.P., Ris, L., and Gabriele, S. (2016) Matrix stiffness modulates formation and activity of neuronal networks of controlled architectures. Biomaterials, 89, 14–24.

50. Beeken, J., Mertens, M., Stas, N., Kessels, S., Aerts, L., Janssen, B., Mussen, F., Pinto, S., Vennekens, R., Rigo, J.-M., Nguyen, L., Brône, B., and Alpizar, Y.A. (2022) Acute inhibition of transient receptor potential vanilloid-type 4 cation channel halts cytoskeletal dynamism in microglia. Glia, 70 (11), 2157–2168.

51. Liot, G., Zala, D., Pla, P., Mottet, G., Piel, M., and Saudou, F. (2013) Mutant Huntingtin Alters Retrograde Transport of TrkB Receptors in Striatal Dendrites. J. Neurosci., 33 (15), 6298–6309.

52. Chavan, D., van de Watering, T.C., Gruca, G., Rector, J.H., Heeck, K., Slaman, M., and Iannuzzi, D. (2012) Ferrule-top nanoindenter: An optomechanical fiber sensor for nanoindentation. Review of Scientific Instruments, 83 (11), 115110.

53. Corne, T., Sieprath, T., and Vandenbussche, J. (2017) Deregulation of focal adhesion formation and cytoskeletal tension due to loss of A-type lamins. Cell Adhesion & Migration, 11, 447–463.

54. Jung, S., Aliberti, J., Graemmel, P., Sunshine, M.J., Kreutzberg, G.W., Sher, A., and Littman, D.R. (2000) Analysis of fractalkine receptor CX(3)CR1 function by targeted deletion and green fluorescent protein reporter gene insertion. Mol Cell Biol, 20 (11), 4106–4114.

